# Ribotin: Automated assembly and phasing of rDNA morphs

**DOI:** 10.1101/2023.09.29.560103

**Authors:** Mikko Rautiainen

## Abstract

The ribosomal DNA (rDNA) arrays are highly repetitive and homogenous regions which exist in all life. Due to their repetitiveness, current assembly methods do not fully assemble the rDNA arrays in humans and many other eukaryotes, and so variation within the rDNA arrays cannot be effectively studied. Here we present the tool ribotin to assemble full length rDNA copies, or *morphs*. Ribotin uses a combination of highly accurate long reads and extremely long nanopore reads to resolve the variation between rDNA morphs. We show that ribotin successfully recovers the most abundant morphs in human and nonhuman genomes. We also find that genome wide consensus sequences of the rDNA arrays frequently produce a mosaic sequence that does not exist in the genome. Ribotin is available on https://github.com/maickrau/ribotin and as a package on bioconda.

## Introduction

The ribosomal DNA (rDNA) arrays are highly repetitive genomic regions coding for ribosomes, the molecular machinery used in translating proteins. Ribosomes are ubiquitous to all life and are necessary for cellular function, and might be a remnant of a hypothetical “RNA world” before life evolved DNA(12). Heterogeneity within ribosomes has been hypothesized to give rise to specialized ribosomes with differential effects on protein synthesis(13). A recent review(19) listed associations between variation in rDNA and cancer, schizophrenia, intellectual disability and aging in humans, but noted that the associations are uncertain due to small sample sizes and limited methods.

Some manual efforts to assemble rDNA arrays in humans and other organisms have been done(4; 17; 18), but to the best of our knowledge there is no automated tool to perform rDNA assembly. Many aspects of the rDNA arrays, from basic structural issues such as the prevalence of inverted sequences and nontandem repeats to more medically relevant issues such as the functional impact of rDNA variation, remain open questions due to the lack of methods to assemble rDNA arrays(19). An automated method to assemble rDNA arrays would enable studies to look at large numbers of human genomes and investigate these questions.

Due to their repetitiveness and homogeneity, the rDNA arrays are the only remaining regions in human genomes which are inaccessible with existing genome assembly methods. Although current state of the art genome assembly tools(1; 11) can assemble nearly everything in human genomes, they do not assemble the rDNA arrays completely outside of a few cases when the rDNA arrays are particularly short.

In humans, the rDNA arrays are located in chromosomes 13, 14, 15, 21 and 22 and are composed of a few hundred copies of an approximately 45kbp repeat unit arranged in tandem repeats. These repeat units are highly similar, with each array typically having dozens of identical or near-identical repeat units. Copies within the same chromosome are more similar than copies in different chromosomes, and there is chromosome specific variation which enables different rDNA copies to be assigned to different chromosomes(4).

The release of the CHM13 telomere-to-telomere genome in 2022(4) provided the first chromosome resolved assembly of the rDNA arrays of one human genome. The rDNA arrays were manually assembled, and due to the difficulty of assembling rDNA arrays, the arrays were filled with model sequence corresponding to the most common repeat units per chromosome duplicated according to their estimated copy counts in arbitrary order. The assembly resolved chromosome specific *morphs*. A morph is the sequence of one complete repeat unit which appears in the rDNA arrays once or multiple times.

Here we present the tool ribotin to automatically assemble rDNA morphs using a combination of long reads. Ribotin requires high accuracy long reads such as PacBio HiFi, and additionally very long reads such as ultralong nanopore reads (ONT) to resolve complete morphs. Since assemblers using HiFi reads sometimes separate the rDNA arrays in different chromosomes due to chromosome specific variation, this information can be used to perform rDNA assembly in a chromosome specific manner. Ribotin has integration with the assembly tool verkko(1) to assemble rDNA morphs per chromosome. Ribotin also has a mode to run without a verkko assembly using only a related reference rDNA sequence.

We test ribotin on human, gorilla and *A. thaliana* genomes, and find that it successfully resolves the morphs of the CHM13 cell line matching previous assembly, resolves rDNA morphs in nonhuman genomes, and in the case of *A. thaliana* discovers nearly all variation in the rDNA arrays. We also perform a small experiment on the HG002 genome showing how the morphs can be used for downstream analysis.

## Methods

Ribotin requires highly accurate long reads, such as PacBio HiFi or Oxford Nanopore Duplex reads, in order to build a genome wide consensus rDNA sequence and detect variation in the rDNA. Additionally, reads long enough to span complete rDNA units are required for resolving complete rDNA morphs.

Figure 1 shows an outline of ribotin. Ribotin first uses the highly accurate long reads to build a graph which represents all variation within the rDNA. Then ultralong ONT reads are aligned to the graph and are used to detect rDNA repeat units. The ONT read paths are then clustered to rDNA morphs, and are used to build a consensus.

**Fig. 1:**
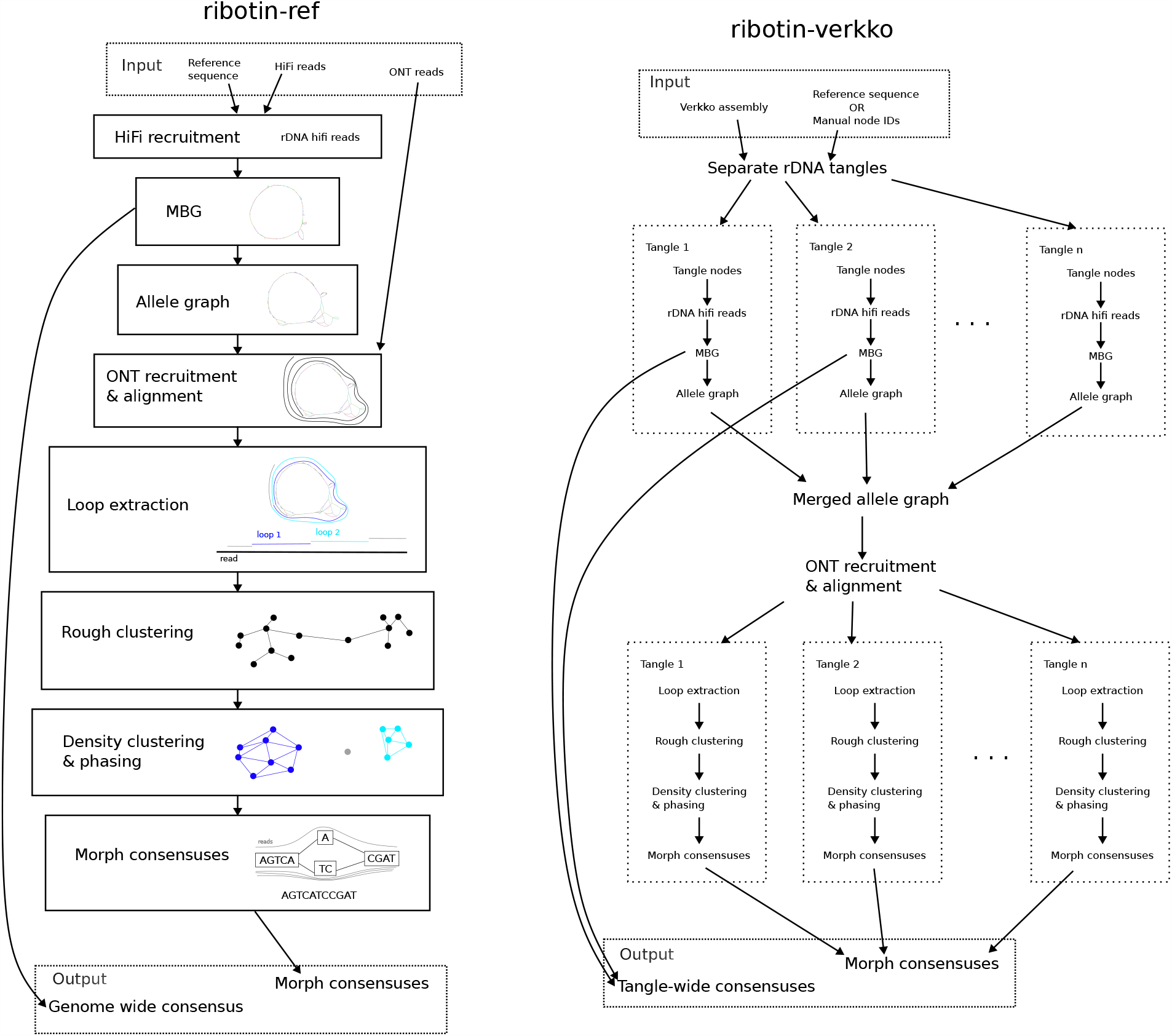
Overview of ribotin. Left: the reference based *ribotin-ref* mode. A reference sequence is used to recruit HiFi reads, which are then assembled with MBG and processed into the allele graph. Then, ONT reads are recruited based on the k-mers of the allele graph and aligned to the allele graph. Loops are extracted from the alignments, which are then clustered to morphs. Finally, a consensus sequence is found for each cluster. Right: the verkko based *ribotin-verkko* mode. A verkko assembly is required, along with either a reference sequence used for detecting rDNA tangles or a manual assignment of node IDs to rDNA tangles. The HiFi reads are assigned to rDNA tangles based on their locations in the verkko assembly. Then, a pipeline similar to ribotin-ref is used per tangle. The ONT reads are recruited and aligned simultaneously to all allele graphs but otherwise the steps are the same as ribotin-ref.

Ribotin has two modes: a *reference-based* mode (*ribotin-ref*), and *verkko-based* mode (*ribotin-verkko*). In ribotin-ref, a reference rDNA sequence is used to recruit HiFi reads. In ribotin-verkko, a reference rDNA sequence is used to detect unresolved rDNA clusters in a verkko(1) assembly, and the hifi reads assigned to those clusters are used. The benefit given by this depends on how well the clusters are separated in the verkko assembly, with the best results when all rDNA arrays are in separate tangles and no benefit when all rDNA arrays are in the same tangle. Additionally, in ribotin-verkko the user may manually select rDNA nodes instead of using a reference sequence to detect them. Ribotin- ref handles all rDNA hifi reads in one go, while ribotin-verkko processes each distinct rDNA tangle separately. The reference sequence does not need to be fully assembled or contain any full length morphs as long as it contains most of the rDNA k-mers present in the genome.

### HiFi Read recruitment

The first step in ribotin-ref is recruiting HiFi rDNA reads. This is done by matching the k-mers of the reference sequence to the reads. The amount of sequence in the read covered by the matching 201-mers is counted, and if the k-mers cover at least 2000 bases, the read is included in the set of rDNA reads.

In ribotin-verkko, the reference sequence is instead matched to nodes in the verkko assembly. 201-mers are matched, and any node with at least 2000 bases covered is a potential rDNA tangle node. Nodes longer than 100kbp are not included as they are considered already well enough resolved. An *rDNA tangle* is a maximal subgraph containing only potential rDNA tangle nodes, with at least 10 nodes, and which contains at least one cycle. Ribotin detects the rDNA tangles using the reference k-mers and graph topology. The user may optionally manually select the rDNA tangles instead of using the default tangle detection. Once the rDNA tangles are found, ribotin recruits HiFi reads which are uniquely assigned to only one tangle, and then runs the next steps separately for each tangle. This step can lose some coverage, since verkko performs graph cleaning which removes low coverage sequences, and discards HiFi reads which are entirely contained within a resolved repeat. We observed that ribotin-verkko typically recruits about 40%-50% fewer HiFi reads than ribotin-ref.

### Graph building and processing

In both ribotin-ref and ribotin-verkko, once the HiFi reads have been recruited, they are processed. First, MBG(3) is used to build a *locally acyclic* de Bruijn graph. We have adjusted MBG to implement a locally acyclic resolution method. In brief, MBG uses the *multiplex de Bruijn graph* algorithm(5) to resolve nodes by increasing *k* locally. In the locally acyclic resolution, this multiplex *k* increase is only done on nodes which are *locally read repetitive*. A node is locally read repetitive with distance *d* if it occurs more than once in a read with a distance of at most *d* between two adjacent occurrences. We set *d* to be one fifth of the estimated morph size provided by the user. This ensures that there are no small cycles within the graph, while preserving a globally cyclic structure to represent the tandemly repeating nature of the rDNA arrays. MBG uses microsatellite error masking(1) to mask away microsatellite indel sequencing errors, which are the second most prevalent sequencing errors in HiFi reads after homopolymer indels. However, some rDNA morphs vary in microsatellite lengths. If the microsatellite length variants are close to other variants, such as SNPs, they will be separated, but if two morphs vary only by microsatellite lengths then this masking artificially homogenizes microsatellite repeats to their most common allele.

Next, ribotin finds a consensus sequence in the graph. The consensus sequence is found with a coverage aware graph walking algorithm. Due to the locally acyclic nature of the graph, any cycle represents one rDNA morph, and the highest coverage cycle is a consensus which picks the highest coverage allele at each bubble. The highest coverage node is selected as the starting point, and then the path is extended to the highest coverage circular path. This produces the *consensus path*, and the sequence of the path is the consensus sequence. This algorithm has previously been used for analysing the CHM13 rDNA arrays(4), and for finding higher order repeats in centromeres(6).

The user may optionally provide a previous reference for orienting the consensus sequence. Matches of unique 101-mers which occur exactly once in both the reference and the consensus are used first to reverse complement the consensus if necessary, and then to rotate it so the two sequences are co-linear. In contrast to the reference used for recruiting reads, the reference used for orienting should contain exactly one full length morph, although it does not need to be from the same individual or species as long as it is similar enough to find 101-mer matches.

Once the consensus sequence is found, variants are detected from the graph. Read subpaths which start at a consensus node, pass through non-consensus nodes and edges, and end at a consensus node are potential variants. Any variant which is supported by at least three reads is outputed. Ribotin further creates an *allele graph*, which contains the consensus nodes and each variant as a path that starts and ends at a consensus node. Nodes which occur in multiple variants are duplicated to one node per variant.

### Morph resolution

Next, the ultralong ONT reads are processed. The first step is recruiting ONT reads by finding all k-mers in the allele graph, and matching those to the ONT reads. 21-mers are used to count the number of base pairs covered by 21-mers, and if the number of matching base pairs is at least half of the consensus sequence length, the ONT read is included. Note that this step uses the k-mers from the allele graph, not from the reference used for recruiting the HiFi reads. The reason for this is that the allele graph is assumed to be more representative of the genome, and therefore better for recruiting. This step does not consider the copy counts or the locations of the k-mers in the graph, only their presence or absence, and it also does not consider their existence elsewhere in the genome.

The ONT reads are then aligned to the allele graph with GraphAligner(2). In ribotin-ref there is only one allele graph, but in ribotin-verkko there is one graph per tangle. These graphs are concatenated into a single file such that GraphAligner will align simultaneously to all of the graphs without allowing a single alignment to cross over between separate graphs. If an ONT read has alignments to multiple graphs, it is discarded, and if it has alignments only to a single graph it is kept for further processing.

The consensus sequence is used to find morph breakpoints in the allele graph. The 201-mers of the consensus sequence are matched to all nodes to find approximate positions for the morph breakpoints. Then the first and last 50bp of the consensus are aligned to the approximate positions to find exact morph breakpoints.

The morph breakpoints are used to extract *loops* from the ONT reads. Whenever the alignment of an ONT read crosses two morph breakpoints, it has completed a single traversal around the allele graph and the path is extracted as a loop. Every loop represents one complete sequence of an rDNA morph, potentially with sequencing errors. All such complete loops are collected from the ONT alignments. Partial loops at the start or end of a read are ignored. We ignore the sequence of the read itself and use the sequence of the path in the graph as the *loop sequence* of a loop. The reason for using the sequences of the graph path instead of the read is that aligning the ONT reads to the graph implicitly performs error correction and enables the reads to be clustered with a more stringent identity threshold.

The loops are then clustered based on sequence similarity. The loops are aligned in an all-vs-all manner using global alignment with the wavefront alignment algorithm(7). Any pair of loops with an edit distance less than the maximum cluster difference *d* (adjustable by user, by default 200 edits corresponding to 99.6% identity) are merged into the same cluster using a union-find data structure. We call these clusters the *rough clusters*. The rough clusters usually contain multiple morphs per cluster, and have the property that any pair of loops between different rough clusters has an edit distance at least *d*. The converse is not true and pairs of loops within the same rough cluster can have any edit distance, including more than *d*.

The rough clusters are then further refined. We cluster the loops by considering each loop a single point, with the distance between points defined by the edit distance of their loop sequences. The ground truth of this clustering problem is that each morph corresponds to one cluster such that there is an unknown number of clusters, the clusters have a different number of points due to their different copy counts, the clusters have a small dense center composed of low error rate points and a large sparse ring of high error rate outliers, and different clusters can be close enough to be indistinguishable if their variation is small compared to the average read error rate.

We use the density based dbscan algorihm(8) for clustering since it does not require the number of clusters to be known, and its parameters can be estimated from the data. The dbscan algorithm requires two parameters: a maximum distance *ϵ*, and a minimum number of points *minPts*, and then it classifies the points into *core points, border points* and *outliers* depending on the number of points close to them. If a point has at least *minPts* points within distance *ϵ*, it is a core point. Any pair of core points within *ϵ* distance of each other belong in the same cluster. If a point is not core but is within *ϵ* of a core point, it is a border point. Otherwise a point is an outlier point. In the basic dbscan algorithm, border points may belong to different clusters depending on the order of iteration and so their assignment can be ambiguous. We detect this case and assign border points to a cluster only if their assignment is unambiguous, that is, all core points within distance *ϵ* are part of the same cluster. Ambiguous border points and outlier points are discarded. We call these clusters the *density clusters*.

The intuition for estimating *ϵ* is that the edit distance histogram between all pairs of loops is multimodal: there is one mode for all loop pairs within the same morph, and one mode for each pair of different morphs. Importantly, since the edit distance between intra-morph loop pairs is composed only of read sequencing errors, the average edit distances of intra-morph loop pairs are the same in all clusters and they are all part of the same mode. We estimate the average intra-morph edit distance by aligning all pairs of loop sequences in the same rough cluster to each others. The peak in the loop pair edit distance histogram is then taken as the average intra-morph edit distance and the *ϵ* parameter. Ribotin further has an option for minimum *ϵ* (by default 5) which limits how low the *ϵ* parameter may be chosen. The *minPts* parameter should be chosen based on the expected number of loops per a single copy morph such that there is a high probability that a single copy morph has at least *minPts* loops. This could be estimated from the genomic coverage and read length distribution, but we use a constant 5 as *minPts* since high copy count morphs will almost certainly have more than 5 loops.

Due to the density based method of dbscan, the same cluster might contain points which are further apart than *ϵ*. This can happen if several highly similar morphs are more similar than *ϵ*, since the cluster will be the transitive closure of the morphs. Therefore the parameter *ϵ* should not be interpreted as a strict limit on the morph resolution, but instead as a guideline on what kind of divergence is resolved, and a single cluster can contain morphs which differ by more than *ϵ*.

Once the loops have been clustered with dbscan, the clusters are then further phased with a heuristic method. The heuristic is based on a simple idea: any sequencing errors will be independent between loops, while valid variation between morphs will co-occur between different loops. If there are two bubbles which split the loops into the same two bipartitions, and both sets in the bipartition are covered by at least 10 loops, then the two sets are separated into their own clusters. This is recursively repeated until the heuristic no longer splits the cluster. This method is very stringent due to requiring an exact bipartition between the two bubbles, and even a single loop misaligning on one of the bubbles will prevent the phasing heuristic from working. It also does not take into account more complicated situations involving multiple variants.

Finally, the consensus for each cluster is generated. A consensus path is generated based on the loop paths in the cluster by taking the most abundant allele at each bubble, and the sequence of the consensus path is outputed as the morph sequence, and the number of loops contained in the cluster as the coverage. These sequences are the final morph consensuses outputed by ribotin.

## Results

We ran ribotin-ref and ribotin-verkko on several genomes using real data. Comparing sets of morphs requires finding similar morphs. When we compare morphs, we align them to each others with minimap2(15) and say that two morphs *match* if they have an alignment with at least 99% identity covering at least 99% of both morphs. We used the parameters ‘-c -x asm5’ for minimap2.

We used ribotin version 1.0, verkko version 1.3.1, MBG version 1.0.15, GraphAligner version 1.0.17b and minimap2 version 2.22- r1101.

### CHM13

The CHM13 assembly(4) is so far the only human assembly with resolved rDNA sequences. We used the existing assembly as the ground truth to evaluate ribotin’s accuracy. We used the same HiFi and ONT data that was used to generate the assembly, consisting of approximately 35x haploid coverage HiFi and 120x ONT.

In ribotin-verkko, chromosomes 13 and 22 were separated due to verkko assembling them in separate tangles. Chromosomes 14, 15 and 21 were assigned to the same tangle. Even though visual inspection with bandage shows that the three rDNA arrays are separate (Supplementary Figure 1), there are short spurious nodes connecting them which causes the tangle detection to consider all three arrays one large tangle. We additionally ran ribotin-verkko in a chromosome specific manner by manually picking the nodes of the five arrays.

In ribotin-ref, the ONT reads successfully phased out chromosome specific morphs. Figure 2 shows how the ribotin- ref morphs match the ribotin-verkko morphs. We observed that ribotin-verkko resulted in more morphs than ribotin-verkko (47 vs 36) due to successfully separating out high similarity low copy count morphs. The *ϵ* parameter chosen by ribotin was highly dependent on how many chromosomes a tangle contained: the two ribotin-verkko tangles which consisted of chromosomes 13 and 22 both had *ϵ* = 5, while the tangle which merged the three remaining chromosomes had *ϵ* = 18. Meanwhile, ribotin-ref which contained all five chromosomes in the same tangle had *ϵ* = 79. The results in Figure 2 support this, showing that the ribotin- ref morphs sometimes match several ribotin-verkko morphs, with the total coverage approximately matching. In a few cases (eg. ribotin-ref IDs 1 and 6 and ribotin-verkko tangle 1 IDs 0 and 2) the morph matchings formed a clique due to resolving morphs which are more than 99% similar and therefore align to each others at 99% similarity. However, there were morphs even within the same chromosome which were less than 99% similar to each others. In the manual chromosome specific mode ribotin chose *ϵ* = 5 for all clusters and produced 52 morphs. This shows that separating the graphs by chromosomes helps with resolving highly similar morphs. The morphs ranged in size from 38kbp to 49kbp, with the average length weighted by coverage being 44683bp.

**Fig. 2:**
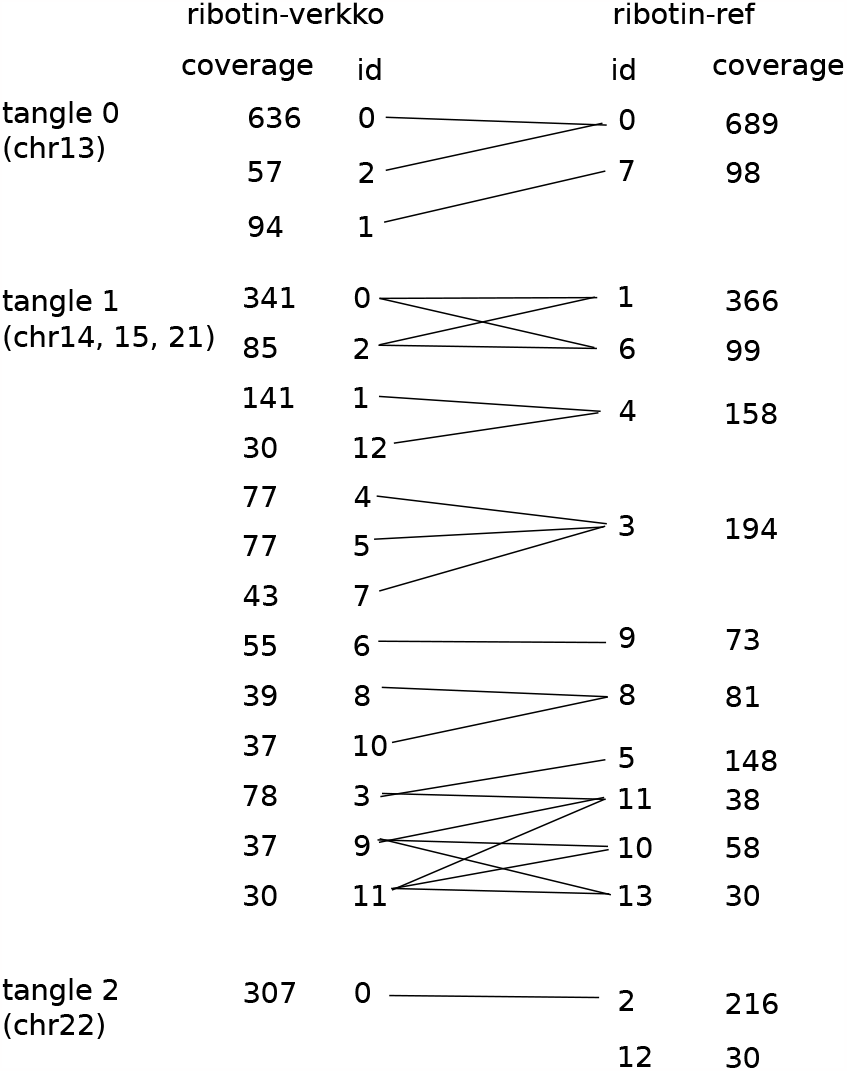
Comparison of ribotin-verkko with automatic rDNA tangle detection and ribotin-ref morphs for CHM13. Morphs with coverage less than 30 are not shown. All matching morphs are connected by lines. The ribotin-verkko morphs are grouped by tangle, and both ribotin-ref and ribotin-verkko morphs are ordered for clarity. Ribotin-ref does not group the morphs in any way and the whitespaces are only for clarity.

We compared the morphs generated by ribotin-ref to the rDNA model sequences resolved by the CHM13 assembly(4). Since the CHM13 assembly rDNA model sequence only includes major morphs, even though the assembly process produced some low copy count morphs((4) Figure S12 panel c), we limit the comparison to the major rDNA morphs present in the assembly. We extracted the major rDNA morphs from the CHM13 assembly, and assigned a copy count based on how many times the exact same morph appears in the assembly. We refer to the previously resolved morphs as CHM13 morphs.

Figure 3 shows how CHM13 morphs and ribotin-ref morphs match each others. All CHM13 morphs are recovered by ribotin. Some of the CHM13 morphs (chr15c, chr21a, chr21b) match two ribotin morphs due to ribotin separating similar but not identical morphs. The highest coverage ribotin morph not found in the CHM13 major morphs was id 7 with coverage 98. Since the CHM13 morphs only included the most abundant morphs, it is expected that some low copy count ribotin morphs are not found in the CHM13 morphs. This shows that the morphs recovered by ribotin match the previous assembly. However, we observed a relatively high average error rate, with matching morphs typically having 100-200 mismatches (0.2% to 0.4% error rate). The error rate was computed from the number of edits of the minimap2 alignment of the highest coverage ribotin morph which matches the CHM13 morph.

**Fig. 3:**
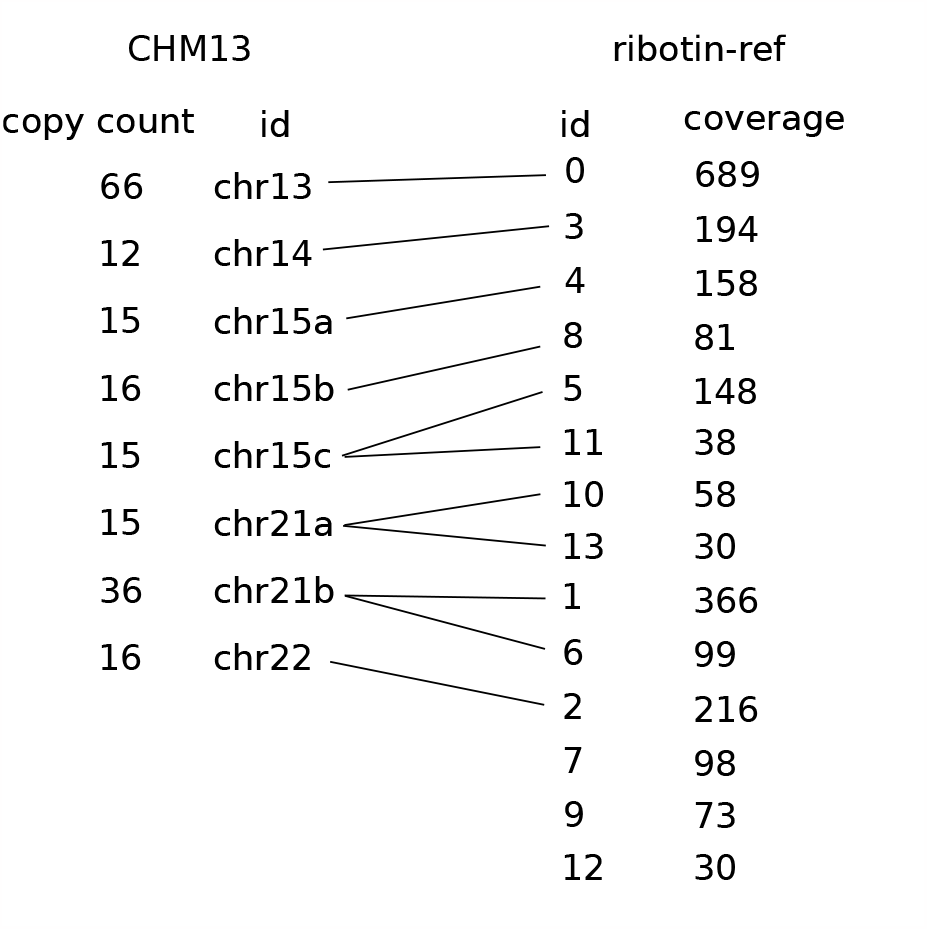
Comparison of the CHM13 major morphs and ribotin-ref morphs. All matching morphs are connected by a line. CHM13 morphs are ordered by ID and ribotin-ref morphs are ordered for clarity. Morphs with coverage less than 30 are not shown.

We also compared the morphs generated by the manual chromosome specific mode to the CHM13 morphs. All CHM13 morphs were recovered. In addition, the error rates were low. The most abundant morph in each chromosome ranged between 0.009% and 0.06% error rate, but the second and third most abundant morphs (chr21b, chr15b, chr15c) had higher error rates from 0.2% to 0.47%. This might be due to MBG collapsing microsatellite variation within each chromosome.

The main difference between the manual chromosome specific mode and the automatic ribotin-ref mode is that the manual mode has a noticably higher consensus accuracy. The manual mode also separated out more low copy count morphs which are highly similar to the abundant morphs, but the high copy count morphs were recovered in both modes. This shows that while separating out the rDNA arrays by chromosome is helpful for getting best results, it is not necessary for recovering the most abundant morphs.

We observed that the genome wide consensus in ribotin- ref only matched to one morph with coverage 30 at 99.5% identity, and didn’t match any of the higher coverage ribotin morphs or the CHM13 morphs. This is due to the presence of chromosome specific variants: all of the chromosomes have some variation present in nearly every copy in the same chromosome but missing from other chromosomes(4), but the single genome wide consensus lacks any of these variants and therefore does not actually correspond to any sequence present in the genome. On the other hand, the chromosome wide consensuses generated by the manual chromosome specific mode, as well as chr13 and chr22 in ribotin-verkko, do match the highest coverage morphs in their chromosomes with very low error rates (0- 4 mismatches corresponding to error rates of 0%-0.009%) and therefore correspond to genomic sequence.

### Human rDNA assembly

We ran ribotin on the human sample HG002. Since this sample does not have a previous rDNA assembly, there is no ground truth to compare to. We compare the consistency of the ribotin-ref morphs to the ribotin-verkko morphs. The sample had 30x HiFi coverage and 60x ONT coverage.

On the HG002 assembly, the CHM13 based reference successfully recruited the relevant HiFi reads and the nodes in the verkko assembly. Ribotin-verkko assigned one rDNA array (chr15 maternal) to its own tangle and all the others in one tangle. Again a visual inspection with bandage (Supplementary Figure 2) suggests that the tangles should be separated, but short nodes connecting the tangles cause them to be merged to the same tangle. In this case the bandage plot is too ambiguous to clearly separate the nine rDNA arrays, so we did not attempt to manually select the rDNA nodes for a chromosome specific assembly.

Ribotin-verkko had two tangles with *ϵ* 96 and 5, while ribotin- ref used *ϵ* = 146. The morphs are roughly consistent between the two modes. One ribotin-verkko morph was split into two in ribotin-ref, and some ribotin-ref morphs were split in ribotin- verkko. This shows again that tangles with fewer chromosomes are better resolved, although the difference is small in this case due to only one array being separated in ribotin-verkko. This shows that the CHM13 reference is similar enough to be used for recruiting rDNA reads in human genomes. The morphs ranged in size from 43kbp to 48kbp, with the average length weighted by coverage being 45239bp. Supplementary Figure 3 shows how the morphs match between ribotin-verkko and ribotin-ref.

We also ran ribotin-ref using a lower coverage dataset with one cell of HiFi from PacBio Sequel II and one cell of ultralong ONT from Promethion, containing 10x HiFi and 35x ONT. We then compared the results to the full coverage ribotin-ref results. Curiously, the lower coverage dataset had an estimated *ϵ* = 75, almost half of the full coverage dataset. Despite this, the higher coverage dataset still resolved the morphs slightly better. The two sets are roughly similar, showing that ribotin can recover abundant morphs even from one cell of HiFi and ONT each. Supplementary Figure 4 shows the results.

We again observed that the genome wide consensus is a poor match to the morphs, with the most similar morph having merely 99.5% identity. The verkko tangle with only one array produced a consensus which does match its highest coverage morph with 2 mismatches (error rate 0.004%).

### Gorilla

We tested ribotin on a gorilla genome to test the performance on a non-human genome. In contrast to the human HG002, we observed that the default CHM13 based reference is not sufficient for recruiting rDNA hifi reads due to dissimilarities between human and gorilla rDNA sequences.

The results using the CHM13 reference misses much variation and most morphs. *ϵ* was estimated at 198, and all loops were clustered into a single morph. To solve this, we used MBG to build an assembly from the whole genome hifi reads, manually picked out the rDNA cluster contigs, and used those as the reference. This resulted in estimated *ϵ* = 95 and produced 6 morphs with coverage ≥ 30. This shows that even in closely related species, there are differences between the rDNA sequences which need to be taken into account.

We also ran ribotin-verkko with the same MBG based reference. Verkko separated the rDNAs into three tangles. Figure 4 shows how the two sets of morphs match. The morphs match between ribotin-ref and ribotin-verkko. The morphs ranged in size from37kbp to 38kbp, with the average length weighted by coverage being 38190bp.

**Fig. 4:**
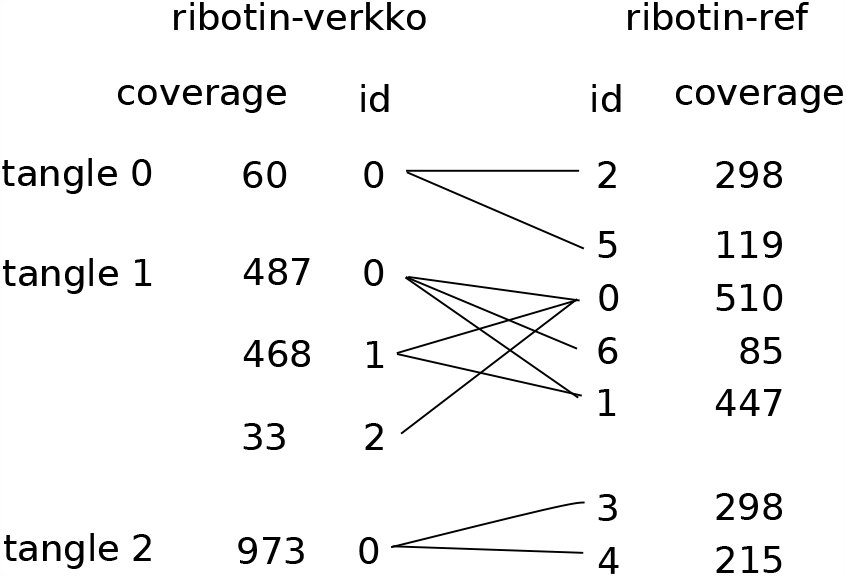
Comparison of ribotin-ref and ribotin-verkko for gorilla. All matching morphs are connected by a line. Morphs with coverage less than 30 are not shown.

The genome wide consensus from ribotin-ref as well as the tangle consensuses from tangles 1 and 2 of ribotin-verkko have an alignment identity of at most 99.5% to a morph. The consensus of tangle 0 matches the highest coverage morph in tangle 0 with one mismatch.

The gorilla morphs were approximately 38kbp long compared to the human 45kbp. Ribotin was given 45000 as the approximate morph size parameter, showing that slight differences between the estimated and real sizes do not matter.

### Arabidopsis thaliana

We also ran ribotin on *A. thaliana* using hifi and ONT data from Wang et al(9). The A. thaliana rDNA is much shorter than human rDNAs at only approximately 10kbp per morph. We again observed that the default CHM13 rDNA reference is not similar enough for recruiting rDNA hifi reads. In fact, using the CHM13 reference, ribotin did not recruit any rDNA hifi reads. We generated a whole genome hifi assembly using MBG, manually located the rDNA tangles by using blast(10) to align the node sequences against the A. thaliana reference, and picked the node sequences in the tangle as the rDNA reference. We used the same node sequences in ribotin-verkko to detect the rDNA tangle. Ribotin-verkko detected only one tangle, resulting in essentially the same pipeline and same results as ribotin-ref.

Since the hifi reads are longer than rDNA morphs (read N50 14936), they can contain entire rDNA morphs, which enables using high accuracy reads to obtain accurate morphs. The whole genome hifi dataset has haploid coverage 170x, with expected 28x coverage of complete single morphs, and expected 0.35x coverage of complete adjacent morphs. We tested this by running ribotin- ref using the HiFi reads as both the hifi reads and ultralong ONT reads, and setting the maximum rough cluster difference to 10 edits and minimum *ϵ* as 1. Ribotin estimated an average within-morph edit distance of 0 which resulted in *ϵ* = 1. This produced 76 distinct morphs with estimated copy count at least one (coverage at least 14), and 11 further morphs with coverage less than half the expected one copy coverage, for a total of 87 morphs. Using expected coverage of 28x and assuming no coverage bias, the morphs have a total estimated copy count of 747. HiFi sequencing has systematically lower coverage in the rDNA arrays(4) and thus the real copy count is likely higher. The most abundant morph had a coverage of 5777, containing slightly over a quarter of the loops. This shows that despite the homogeneity of the rDNA arrays, this genome did not have a single morph which would account for the majority of the rDNA sequence. The morphs ranged in size from 9kbp to 12kbp, with the average length weighted by coverage being 10563bp.

The genome wide consensus sequence matched the fourth most abundant morph (coverage 845) with two mismatches. In addition, three other morphs matched the genome wide consensus with less than 10 mismatches. The four morphs had a total coverage of 1787, or approximately 8.5% of the loops. Curiously, the exact consensus sequence did not exist among the morphs and the most abundant morph had 691 edits (divergence 6.2%) to the consensus sequence.

We believe these morphs contain nearly all rDNA variation that occurs in this A. thaliana individual as well as their relative abundances. Due to ribotin’s choice of *ϵ* = 1, morphs which differ by one edit are collapsed together. We cannot rule out the possibility that low coverage morphs which differ by a single small variant are collapsed into other morphs, but morphs with two or more variants, or a single variant larger than 5bp, are very likely separated. Although the HiFi reads are long enough to span individual rDNA morphs, they rarely span two or more morphs, and so this approach did not enable finding the exact order of the morphs. Even longer high accuracy reads such as ONT duplex might enable a similar approach to resolve the order of the morphs as well.

### HG002 morph pangenome

To show how the morphs might be used for studying variation within the rDNA arrays, we built a *morph pangenome* out of the HG002 ribotin-ref morphs. We used minimap2(15) to align all the morphs against each others, then ran seqwish(16) to build a pangenome graph out of them. This produced a graph with 2686 nodes and 4004 edges. Figure 5 shows a bandage plot of the graph. We observed that there was a considerable amount of variation even within just this one genome. The graph had 764 bubbles, each corresponding to one variant site, of which 386 had more than two alleles. The longest stretch without any variants was only 415 base pairs long.

**Fig. 5:**
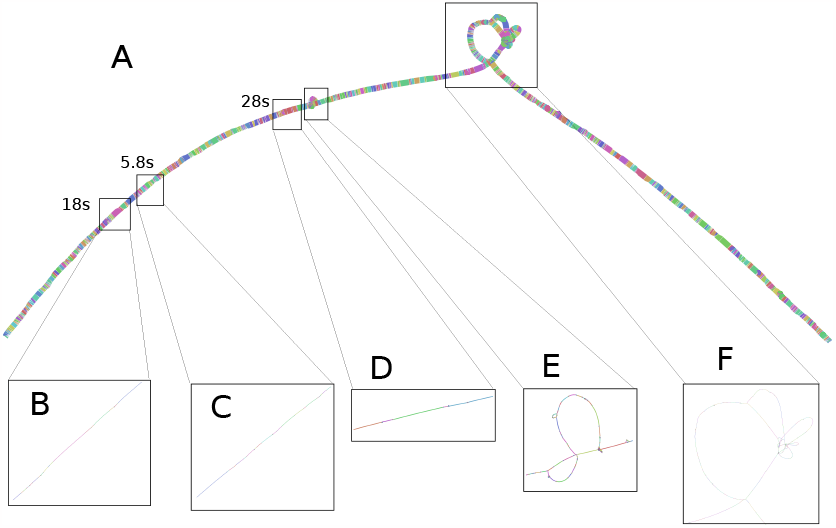
Bandage plot of the morph pangenome of HG002. A: an overview of the entire graph. B: the 18s locus. C: the 5.8s locus. D: the 28s locus. E, F: regions containing structural variation.

We used GraphAligner to align the genome wide HG002 consensus sequence to the graph, and the sequences of the 18s, 5.8s and 28s genes from the KY962518.1 rDNA reference sequence. The genome wide consensus sequence (which was not used in building the graph) aligned to the graph with zero mismatches as expected, taking a path which is a mosaic of different morphs without exactly matching any of the morphs fully. Since the KY962518.1 rDNA reference sequence is from a different individual, the 18s and 5.8s genes aligned to the graph with 1 and 2 mismatches respectively, showing that it contains variation not present in HG002. However, the alignments with 1 and 2 mismatches were alignments to mosaics of the HG002 morphs. Using minimap2 to align the 18s and 5.8s genes to the morphs intead of the graph, the highest identity alignments had 8 and 4 mismatches respectively. The 28s gene aligned with zero mismatches both to the graph and to multiple morphs, including the most abundant morph.

The 18s locus (Figure 5 B) had 23 variants, composed of 9 biallelic SNPs, 9 biallelic indels, and 5 sites with multiple alleles. The 5.8s locus (Figure 5 C) had 26 variants, with 11 biallelic SNPs, 11 biallelic indels and 4 sites with multiple alleles. The 28s locus (Figure 5 D) had 6 variants, with 3 biallelic SNPs and 3 sites with multiple alleles. There were also regions showing structural variation between the morphs (Figure 5 E and F). Although this is basically a toy experiment, it demonstrates how the morphs could be applied for downstream analysis.

### Runtime

Table 1 shows the runtime of ribotin with each dataset and mode. Ribotin was given 8 threads in all runs. The runtime and peak memory was measured by the slurm scheduler’s “seff” command. We see that the runtimes are modest and memory use is low, enabling ribotin to be easily ran on a large number of samples. Human samples can be processed in a few hours with just several Gb of memory. Curiously, the shorter *A. thaliana* genome required much more memory than human. This might be due to the very high coverage of the *A. thaliana* dataset. Note that the runtime only includes running ribotin and does not include the computational cost of running verkko, which is larger than ribotin by multiple orders of magnitude.

**Table 1.** Runtime of ribotin measured by *seff*.

| Sample | Method | CPU-time<br>(h:m:s) | Wall time<br>(h:m:s) | RAM<br>(Gb) |
| --- | --- | --- | --- | --- |
| CHM13 | ribotin-ref | 06:58:56 | 03:14:03 | 1.40 |
|  | ribotin-verkko | 06:19:58 | 02:52:53 | 2.08 |
|  | ribotin-verkko (manual) | 08:20:36 | 03:19:35 | 2.17 |
| HG002 | ribotin-ref | 04:18:24 | 02:30:54 | 2.64 |
|  | ribotin-verkko | 02:28:51 | 01:15:28 | 2.92 |
|  | ribotin-ref (low coverage) | 01:46:41 | 00:55:06 | 0.63 |
| Gorilla | ribotin-ref | 35:45:58 | 05:40:31 | 0.98 |
|  | ribotin-verkko | 03:20:45 | 01:01:40 | 2.22 |
| *A. thaliana* | ribotin-ref | 41:21:35 | 33:13:34 | 40.42 |
|  | ribotin-verkko | 32:58:31 | 26:00:38 | 18.34 |
|  | ribotin-ref (HiFi) | 06:20:32 | 02:59:34 | 39.73 |

## Discussion

Ribotin enables easy, automated rDNA morph assembly for humans. This can be used to generate model rDNA arrays for telomere-to-telomere assembly efforts, since large rDNA arrays are not assembled by current genome assemblers. Our recommendation for telomere-to-telomere assembly efforts is to use ribotin-verkko with manually chosen rDNA arrays in order to generate the most accurate and complete results. The downside of this is that it involves manual curation. For projects involving more than one genome, we suggest using either the automatic mode of ribotin-verkko or ribotin-ref, depending on if the pipeline already involves running verkko. If the project is about nonhumans, we additionally suggest creating a species-specific reference from one of the samples by performing de novo whole genome assembly and extracting the contigs containing rDNA sequence.

In some genomes such as some plants, the morph size may be short enough to be spanned by HiFi reads. In this case ribotin can resolve nearly all variation within the rDNA arrays.

Currently, ribotin has a limitation that copy counts are not estimated. Although the relative abundances of the morphs are counted based on the ONT coverage, this must be translated to copy counts by the user. Another limitation is that the order of the rDNA morphs is not resolved. This is unlikely to be possible with current sequencing technologies due to the size, repetitiveness and homogeneity of the rDNA arrays. Future sequencing technologies combining very long read lengths with high sequencing accuracy might make it possible to resolve the order of the rDNA morphs, and therefore assemble large rDNA arrays completely. Ribotin also does not assign the morphs to chromosomes, and this must be done manually by the user.

The lack of existing assemblies of complete rDNA morphs makes it difficult to evaluate the accuracy of the results. We have shown that ribotin successfully recovered all major morphs of the CHM13 genome, the so far only human genome assembly which has resolved chromosome specific morphs. However, the other experiments only compared the consistency of ribotin-ref to ribotin-verkko due to a lack of ground truth.

The resolution of rDNA morphs depends on how well the rDNA arrays are separated into different tangles. In the CHM13 experiment, manually selecting the five rDNA arrays from the verkko assembly resolved different morphs down to an accuracy of 5 edits, meaning that morphs more similar than 5 edits are not distinguished. Meanwhile, in ribotin-ref which processes all of the rDNA arrays with the same graph, the accuracy was 79 edits. Although the highly abundant chromosome specific morphs were recovered in both cases, the higher accuracy allowed distinguishing several low copy count morphs which are highly similar to the abundant morphs.

Despite the high homogeneity of rDNA arrays, there was noticable variation between the morphs in all of the genomes we tested. In all of the genomes there were morphs which are less than 99% identical to each others. The lengths of the assembled morphs also varied. In CHM13, the shortest morph was over 10kbp shorter than the longest one. On the other hand, in HG002 the difference was about 5kbp. The gorilla morphs were relatively more homogenous, with only 1.2kbp length difference. Even the *A. thaliana* morphs, despite their short average size of around 10kbp, varied by 3kbp between the shortest and longest.

Due to chromosome specific variation between rDNA arrays, a single genome wide rDNA consensus might result in a sequence which does not exist in the genome at all. Even though the individual alleles of the consensus sequence at each variant position do exist in the genome, the combination of alleles chosen by the consensus might not. We observed this to be the case in both human genomes we tested as well as the gorilla, where the consensus had at best 99.5% alignment identity with any of the morphs. On the other hand, in *A. thaliana* the consensus exists in the genome although with a relatively low copy count. Based on this we suggest that caution should be used when analysing genome wide rDNA consensuses. This highlights the importance of using resolved, complete rDNA morphs instead of a single genome wide rDNA consensus.

## Conclusion

We have presented ribotin, a tool for assembling rDNA morphs. Ribotin can automatically assemble the most abundant morphs in human genomes, and with a little manual processing, nonhuman genomes as well. Ribotin can enable genomic studies to examine the variation within the rDNA arrays which has so far been difficult to analyze due to their high repetitiveness.

## Author contributions statement

## Acknowledgments

The authors wish to acknowledge CSC – IT Center for Science, Finland, for computational resources. We are grateful to Samuli Ripatti and Aarno Palotie for feedback on a manuscript draft.

## References

1. Mikko Rautiainen, Sergey Nurk, Brian P Walenz, Glennis A Logsdon, David Porubsky, Arang Rhie, Evan E Eichler, Adam M Phillippy and Sergey Koren. Telomere-to-telomere assembly of diploid chromosomes with Verkko Nature Biotechnology, 2023.

2. Mikko Rautiainen and Tobias Marschall. GraphAligner: rapid and versatile sequence-to-graph alignment Genome Biology, 2020.

3. Mikko Rautiainen and Tobias Marschall. MBG: Minimizer-based sparse de Bruijn Graph construction Bioinformatics, 2020.

4. Sergey Nurk et al. The complete sequence of a human genome. Science, 2022.

5. Anton Bankevich, Andrey V Bzikadze, Mikhail Kolmogorov, Dmitry Antipov and Pavel A Pevzner. Multiplex de Bruijn graphs enable genome assembly from long, high-fidelity reads. Nature Biotechnology, 2022.

6. Yujie Zhang, Justin Chu, Haoyu Cheng and Heng Li. De novo reconstruction of satellite repeat units from sequence data. arXiv, 2023.

7. Santiago Marco-Sola, Juan Carlos Moure, Miquel Moreto, Antonio Espinosa. Fast gap-affine pairwise alignment using the wavefront algorithm. Bioinformatics, 2020.

8. Martin Ester, Hans-Peter Kriegel, J”org Sander and Xiaowei Xu. A Density-Based Algorithm for Discovering Clusters in Large Spatial Databases with Noise. Proceedings of the Second International Conference on Knowledge Discovery and Data Mining, 1996.

9. Bo Wang, Xiaofei Yang, Yanyan Jia, Yu Xu, Peng Jia, Ningxin Dang, Songbo Wang, Tun Xu, Xixi Zhao, Shenghan Gao, Quanbin Dong and Kai Ye. High-quality Arabidopsis thaliana Genome Assembly with Nanopore and HiFi Long Reads. Genomics, proteomics & bioinformatics, 2022.

10. Stephen F. Altschul, Warren Gish, Webb Miller, Eugene W. Myers and David J. Lipman. Basic local alignment search tool. Journal of molecular biology, 1990.

11. Haoyu Cheng, Mobin Asri, Julian Lucas, Sergey Koren and Heng Li. Scalable telomere-to-telomere assembly for diploid and polyploid genomes with double graph. arXiv, 2023.

12. Thomas R. Cech. The RNA Worlds in Context. Cold Spring Harb Perspect Biol, 2012.

13. Shifeng Xue and Maria Barna. Specialized ribosomes: a new frontier in gene regulation and organismal biology. Nat Rev Mol Cell Biol, 2012.

14. Ryan R. Wick, Mark B. Schultz, Justin Zobel and Kathryn E. Holt. Bandage: interactive visualization of de novo genome assemblies. Bioinformatics, 2015.

15. Heng Li. Minimap2: pairwise alignment for nucleotide sequences. Bioinformatics, 2018.

16. Erik Garrison and Andrea Guarracino. Unbiased pangenome graphs. Bioinformatics, 2023.

17. Qiutao Ding, Runsheng Li, Xiaoliang Ren, Lu-yan Chan, Vincy W. S. Ho, Dongying Xie, Pohao Ye and Zhongying Zhao. Genomic architecture of 5S rDNA cluster and its variations within and between species. BMC Genomics, 2022.

18. Yutaro Hori, Akira Shimamoto and Takehiko Kobayashi. The human ribosomal DNA array is composed of highly homogenized tandem clusters. Genome Research, 2021.

19. Ashley N. Hall, Elizabeth Morton and Christine Queitsch, First discovered, long out of sight, finally visible: ribosomal DNA. Trends in Genetics, 2022.

