## Supplementary Material for "Ribotin: Automated assembly and phasing of rDNA morphs"

### Supplementary Material: Ribotin: Automated assembly and phasing of rDNA morphs

#### A. Supplementary figures

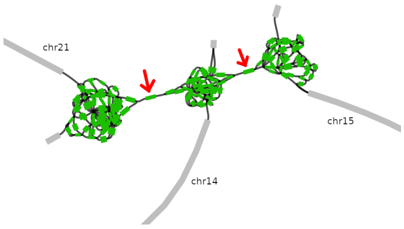


Figure 1: Bandage (Wick *et al.*, 2015) plot of the verkko assembly of CHM13, zoomed in on the rDNA arrays of chromosomes 14, 15, 21. The green nodes were classified as rDNA tangle nodes by ribotin. The two small nodes connecting the tangles (red arrows) caused all three arrays to be assigned to the same tangle.

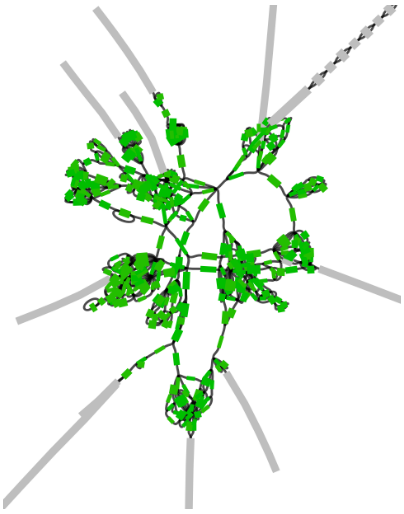

Figure 2: Bandage (Wick *et al.*, 2015) plot of the verkko assembly of HG002, zoomed in on the rDNA arrays. The green nodes were classified as rDNA tangle nodes by ribotin. Visually there appear to be several arrays which were all assigned to the same tangle.

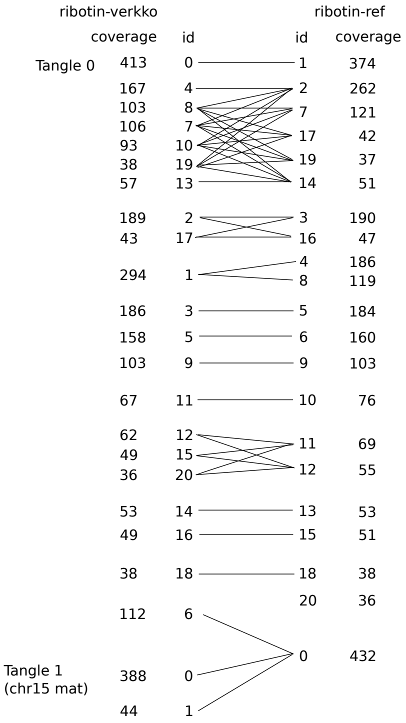


Figure 3: Comparison of ribotin-verkko with automatic rDNA tangle detection and ribotin-ref morphs for HG002. Morphs with coverage less than 30 are not shown. All matching morphs are connected by lines. The ribotin-verkko morphs are grouped by tangle, and both ribotin-ref and ribotin-verkko morphs are ordered for clarity.

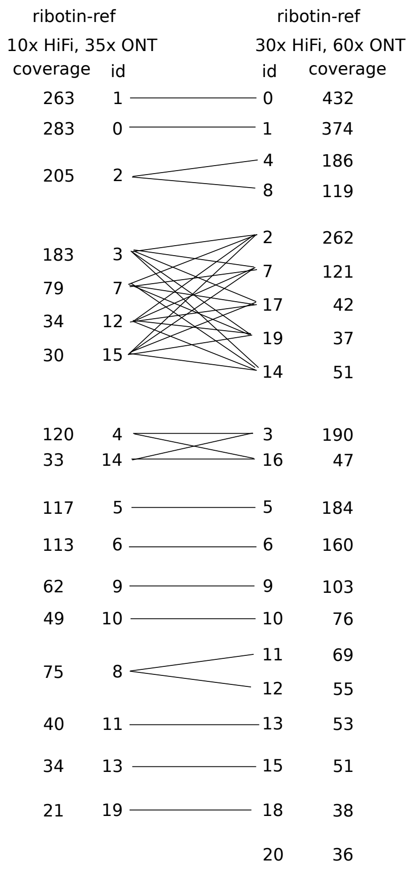


Figure 4: Comparison of ribotin-ref with high and low coverage for HG002. Morphs with coverage less than 30 are not shown. All matching morphs are connected by lines. The morphs are ordered for clarity.

#### References

Wick, R. R., Schultz, M. B., Zobel, J., and Holt, K. E. (2015). Bandage: interactive visualization of de novo genome assemblies. *Bioinformatics*.
